# Optic Atrophy-associated TMEM126A is an assembly factor for the ND4-module of Mitochondrial Complex I

**DOI:** 10.1101/2020.09.18.303255

**Authors:** Luke E. Formosa, Boris Reljic, Alice J. Sharpe, Linden Muellner-Wong, David A. Stroud, Michael T. Ryan

**Affiliations:** Department of Biochemistry and Molecular Biology, Monash Biomedicine Discovery Institute, Monash University, 3800, Melbourne, Australia; Department of Biochemistry and Molecular Biology, The Bio21 Institute, The University of Melbourne, 3000, Melbourne, Australia

## Abstract

Mitochondrial disease is a debilitating condition with a diverse genetic aetiology. Here, we report that TMEM126A, a protein that is mutated in patients with autosomal recessive optic atrophy, participates directly in the assembly of mitochondrial complex I. Using a combination of genome editing, interaction studies and quantitative proteomics, we find that loss of TMEM126A results in an isolated complex I deficiency and that TMEM126A interacts with a number of complex I subunits and assembly factors. Pulse-labelling interaction studies reveal that TMEM126A associates with the newly synthesised mtDNA-encoded ND4 subunit of complex I. Our findings indicate that TMEM126A is involved in the assembly of the ND4 distal membrane module of complex I. Importantly, we clarify that the function of TMEM126A is distinct from its paralogue TMEM126B, which acts in assembly of the ND2-module of complex I, helping to explain the differences in disease aetiology observed between these two genes.

## Introduction

Mitochondria are hubs of metabolic and biosynthetic activity, and also play important roles in a variety of cellular processes that include antiviral signalling, calcium handling and cell death (Spinelli and Haigis, 2018). Mutations in at least 289 genes that encode mitochondrial proteins have a negative effect on mitochondrial homeostasis and are associated with mitochondrial disease (Frazier et al., 2019). A large proportion of these mitochondrial proteins have a direct role in energy generation as they are required as structural subunits or assembly factors for the oxidative phosphorylation (OXPHOS) system (Signes and Fernandez-Vizarra, 2018). Additional gene products that are mutated in mitochondrial disease have various roles including the maintenance and expression of mitochondrial DNA (mtDNA), assembly of iron-sulfur clusters, lipid metabolism, mitochondrial dynamics and protein import/processing that are secondary to energy generation processes (Gorman et al., 2016). However, in many cases, it is unclear what the molecular role of some mitochondrial disease-associated genes are, making clinical interventions difficult.

One such example is the role of TMEM126A in mitochondrial function. A homozygous nonsense mutation in the gene encoding TMEM126A (p. Arg55X) was first identified in individuals with autosomal-recessive optic atrophy (AROA) (Hanein et al., 2009). Since then, other patients with AROA have been identified with the same mutation in the *TMEM126A* gene (Desir et al., 2012; Meyer et al., 2010). The proximal geography of the individuals identified (Morocco and Tunisia), together with similarities in microsatellite analysis at that locus suggest a founder effect may be responsible for this mutation (Hanein et al., 2009). More recently, individuals with additional novel missense mutations in *TMEM126A* have been identified with AROA (Kloth et al., 2019; La Morgia et al., 2019). Patients identified with mutations in *TMEM126A* generally present with bilateral deficiency in visual acuity with onset from infancy to 6 years of age, optic nerve pallor and central scotoma (Hanein et al., 2009). Retinal ganglion cells are particularly susceptible to mitochondrial dysfunction as the intraocular length of the axons that form the optic nerve remain unmyelinated, and so are highly dependent on ATP to propagate the action potential to the visual cortex (Carelli et al., 2009). In addition to AROA, some patients also presented with moderate hypertrophic cardiomyopathy and mild hearing loss (Hanein et al., 2009).

TMEM126A has been reported to be a mitochondrial inner membrane protein, that is enriched in the cristae (Hanein et al., 2013), however its molecular function is unclear. While TMEM126A is metazoan specific, its paralog TMEM126B is found in mammals following a gene duplication event (Elurbe and Huynen, 2016). TMEM126B is essential for the biogenesis of respiratory chain complex I as a membrane-integrated assembly factor and central component of the mitochondrial complex I intermediate assembly (MCIA) complex (Andrews et al., 2013; Formosa et al., 2020; Heide et al., 2012). Complex I is assembled via the integration of 45 subunits into a number of intermediate modules and at least 15 assembly factors that come together in the mitochondrial inner membrane (Formosa et al., 2018; Guerrero-Castillo et al., 2017; Lazarou et al., 2007; Sánchez-Caballero et al., 2016a; Stroud et al., 2016; Ugalde et al., 2004). The MCIA complex is involved in assembly of the ND2 membrane module (Formosa et al., 2020; Guerrero-Castillo et al., 2017). While it has been suggested that TMEM126A may play a compensatory role in patient cells with mutations in TMEM126B (Sánchez-Caballero et al., 2016b), the clinical presentation of the two markedly differ (AROA vs late onset myopathy) (Alston et al., 2016). Nevertheless, a role for TMEM126A in complex I assembly may be implicated given that it was found enriched with the complex I assembly factors DMAC1 (Stroud et al., 2016), TMEM186 and COA1 (Formosa et al., 2020), as well as with the general mitochondrial insertase OXA1L (Thompson et al., 2018).

Given that mutations in mitochondrially encoded complex I subunits can result in a range of diseases including Leber’s Hereditary Optic Neuropathy (LHON) (De Vries et al., 1996; Lin et al., 2018), we sought to determine if TMEM126A plays a direct or indirect role in the assembly of mitochondrial OXPHOS complexes, particularly complex I. Complete loss of TMEM126A results in isolated complex I deficiency, with decreased levels of the fully assembled enzyme as well individual complex I subunits. Furthermore, we show that TMEM126A interacts with complex I subunits and assembly factors under steady-state conditions and enriches with the newly translated mitochondrially-encoded ND4 subunit. Importantly, we establish that TMEM126A does not play a compensatory role to TMEM126B, but rather highlight that it functions in the biogenesis of a distinct assembly module of complex I.

## Results

### Complex I deficiency in TMEM126A knockout cells

In order to investigate the role of TMEM126A in mitochondrial function, we generated knockout (T126A^KO^) cell lines using genome editing. SDS-PAGE and subsequent western blot analysis of isolated mitochondria revealed the absence of TMEM126A from T126A^KO^ cells (**Fig. 1A**). Sequencing of alleles also confirmed the presence of indels that disrupt the open reading frame (**Table S1**). Next, to determine how the loss of TMEM126A affected the mitochondrial proteome, unbiased SILAC quantitative proteomics analysis was performed (**Fig. 1B; Table S2**). Complex I subunits were clearly reduced in the T126A^KO^ mitochondria relative to the controls (**Fig. 1B**). Using the proteomic data from T126A^KO^ cells, a topographical proteomic heat-map was fitted onto the molecular structure of human complex I (Gu et al., 2016). Subunits belonging to distinct assembly modules of complex I – namely the ND2, ND4 - were significantly reduced relative to control mitochondria (**Fig. 1C**).

**Figure 1:**
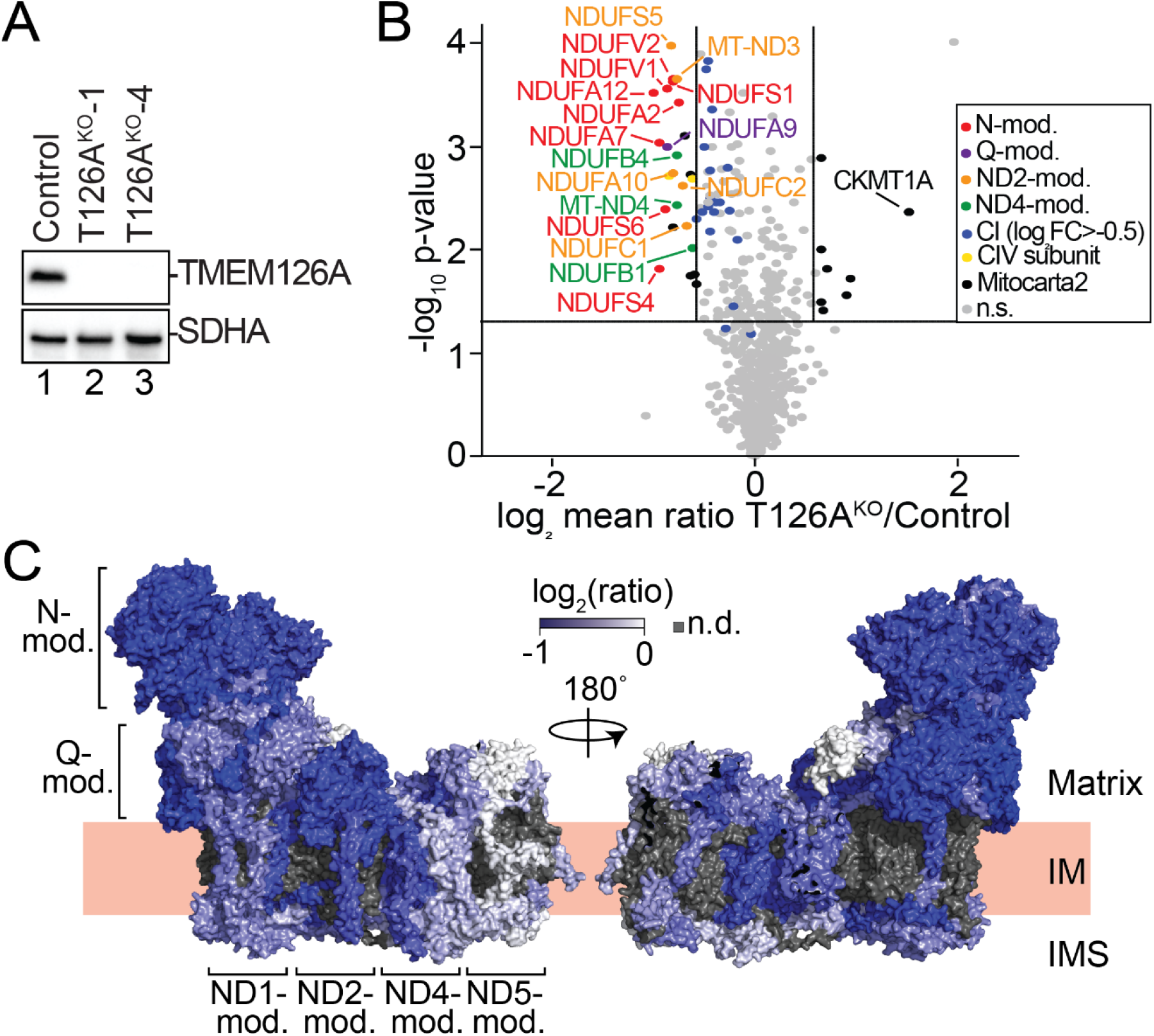
Complex I subunits reduced in T126A^KO^ mitochondria. **(A)** Isolated mitochondria from control and T126A^KO^ cell lines were subjected to SDS-PAGE and immunoblot analysis. SDHA was used as a loading control. **(B)** Volcano plot of protein abundance in T126A^KO^ mitochondria relative to control. SILAC ratios are log_2_ transformed and plotted against −log_10_(p-value). n=3. Horizontal line indicates *p*=0.05; vertical lines indicate 1.5-fold change. **(C)** Topographical heatmap of subunit ratios were fitted to complex I (PDB: 5XTH) (Gu et al., 2016). Grey regions represent subunits that were not reliably detected.

Loss of these subunits correlate with defects in the biogenesis of the membrane arm of complex I (Stroud et al., 2016). The N-module is commonly reduced in all cases where complex I assembly is perturbed (Stroud et al., 2016). In contrast, the most upregulated protein in T126A^KO^ mitochondria was the U-type Creatine Kinase CKMT1A, an enzyme required for maintaining energy homeostasis and buffering ATP levels in cells (Schlattner et al., 2006).

### Loss of TMEM126A leads to an isolated complex I defect

Given the observed decrease in complex I subunits by SILAC proteomics, we isolated mitochondria from control and T126A^KO^ cells and assessed complex I assembly by BN-PAGE and western blotting (**Fig. 2A**). Solubilisation of mitochondrial membranes in the mild detergent digitonin maintains complex I in its supercomplex form with complexes III and IV, while the harsher detergent Triton X-100 dissociates the supercomplex into the holo-complex forms (McKenzie et al., 2007). Immunoblotting for the complex I subunit NDUFB8 revealed a reduction in the levels of the supercomplex and in the complex I holoenzyme in comparison to control mitochondria (**Fig. 2A**). Quantification of the NDUFB8 signal showed that the supercomplex was reduced by ~50% of control levels, while the levels of holo-complex I were reduced to ~30% to that of control mitochondria (**Fig. 2B**). In addition, a faster migrating species at ~440 kDa following digitonin solubilization and ~400 kDa following Triton X-100 solubilization was also apparent (indicated by *; **Fig. 2A**). Given that the NDUFB8 subunit is located at the distal end of the complex I membrane arm (Fiedorczuk et al., 2016; Zhu et al., 2016), this subcomplex may represent an accumulation or breakdown product of this region in the absence of proper complex I assembly. Besides the complex I supercomplexes, OXPHOS complexes III, IV or V were not changed in T126A^KO^ mitochondria (**Fig. 2C**; note levels of holo-complexes III and IV in lanes 4-6 and 10-12 respectively), consistent with proteomic analysis.

**Figure 2:**
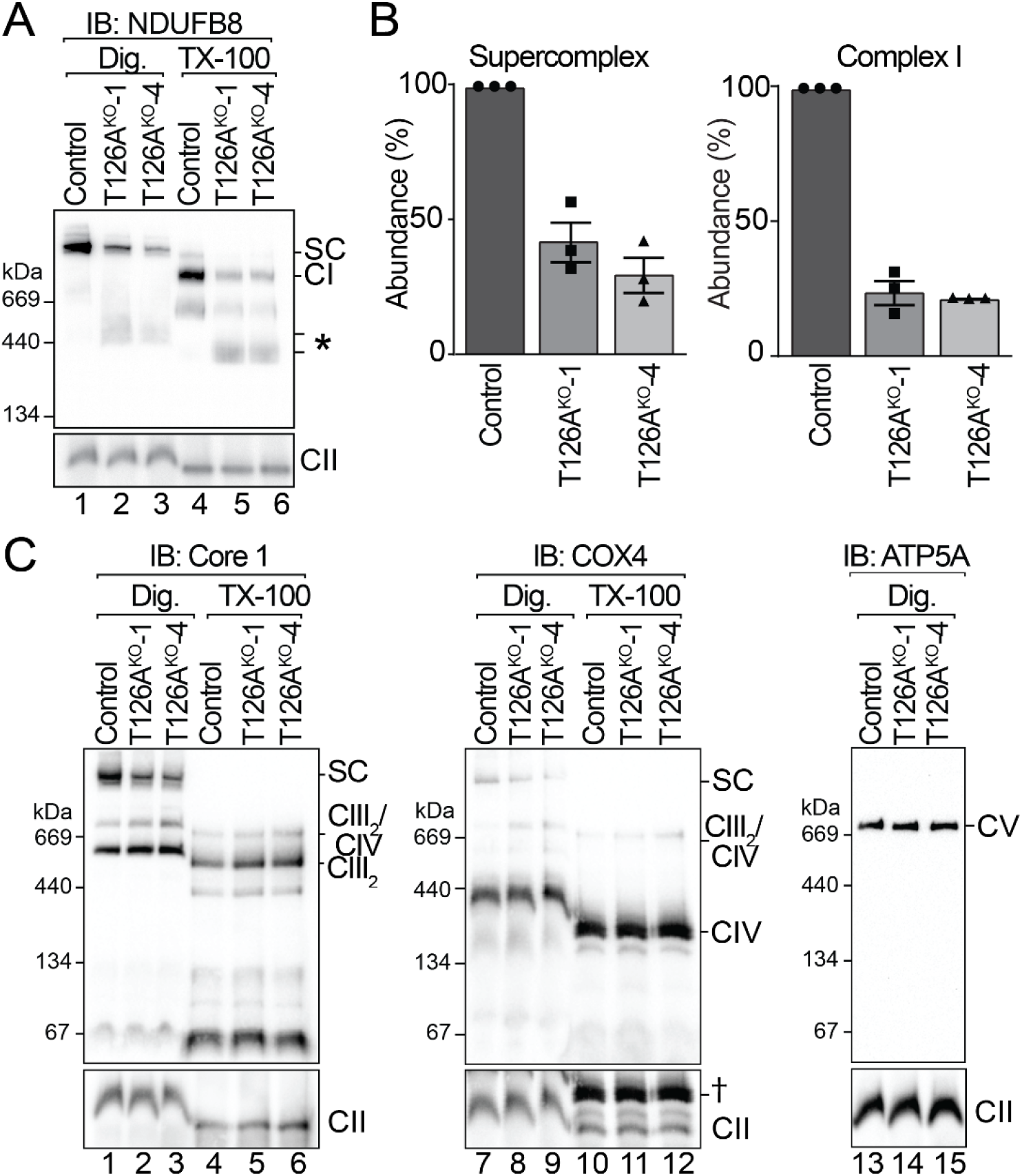
T126A^KO^ cells have an isolated complex I deficiency. **(A)** Mitochondria were isolated from control and T126A^KO^ cell and subjected to BN-PAGE and western blotting for complex I (NDUFB8) and complex II (SDHA) used as a control. **(B)** The relative abundance of fully assembled supercomplex and complex I was quantified (mean ± SEM, n=3). **(C)** Isolated mitochondria subjected to BN-PAGE and western blotting for complexes III (Core 1), IV (COX4) and V (ATP5A). † residual COX4 signal following reprobe.

### TMEM126A associates with complex I subunits, assembly factors and newly-synthesised ND4

Next, we complemented the knockout cells with a construct expressing TMEM126A with a C-terminal Flag epitope tag (TMEM126A^Flag^) and demonstrated rescue of the complex I assembly defect (**Fig. 3A**). We subsequently performed affinity-enrichment and label-free quantitative (LFQ) mass spectrometry analysis to determine TMEM126A interacting partners. TMEM126A^Flag^ was able to enrich all accessory subunits of the ND4-module (NDUFB1, NDUFB5, NDUFB10 and NDUFB11) as well some subunits of the Q- (NDUFS2), ND1- (NDUFA8, NDUFA13 and NDUFA3), ND2- (NDUFA10) and ND5- (NDUFB6, NDUFB8 and NDUFB9) modules (**Fig. 3B; Table S3**). Furthermore, TMEM126A^Flag^ enriched a number of complex I assembly factors including the MCIA complex factors ECSIT and ACAD9, the ND1-module assembly factor TIMMDC1, plus the Q-module assembly factors NDUFAF3 and NDUFAF4. TMEM70, an assembly factor recently implicated in the assembly of the complex I ND4-module was also enriched (Guerrero-Castillo et al., 2017; Sánchez-Caballero et al., 2020). In agreement with previous studies (Thompson et al., 2018), we also observed that TMEM126A^Flag^ was able to enrich the general membrane insertase for mtDNA encoded proteins, OXA1L. Defects in complex I assembly lead to the accumulation of assembly intermediates consisting of assembly factors and complex I modules. Neither the major MCIA complex at ~450 kDa (involved in ND2-module assembly; **Fig. 3C**, ◊) nor the TIMMDC1 complexes at ~400/440 kDa (involved in ND1-module assembly; **Fig. 3C**, §) were altered in T126A^KO^ mitochondria. However, a higher molecular-weight doublet species of ~680/720 kDa (containing both MCIA and TIMMDC1; **Fig. 3C**, #) representing intermediates further along the assembly pathway, was reduced in T126A^KO^ relative to control mitochondria (**Fig. 3C**). These results suggest that while TMEM126A may not participate in complex I assembly directly with the MCIA or TIMMDC1 assembly complexes, loss of TMEM126A impairs later stages of the assembly pathway involving these machineries.

**Figure 3:**
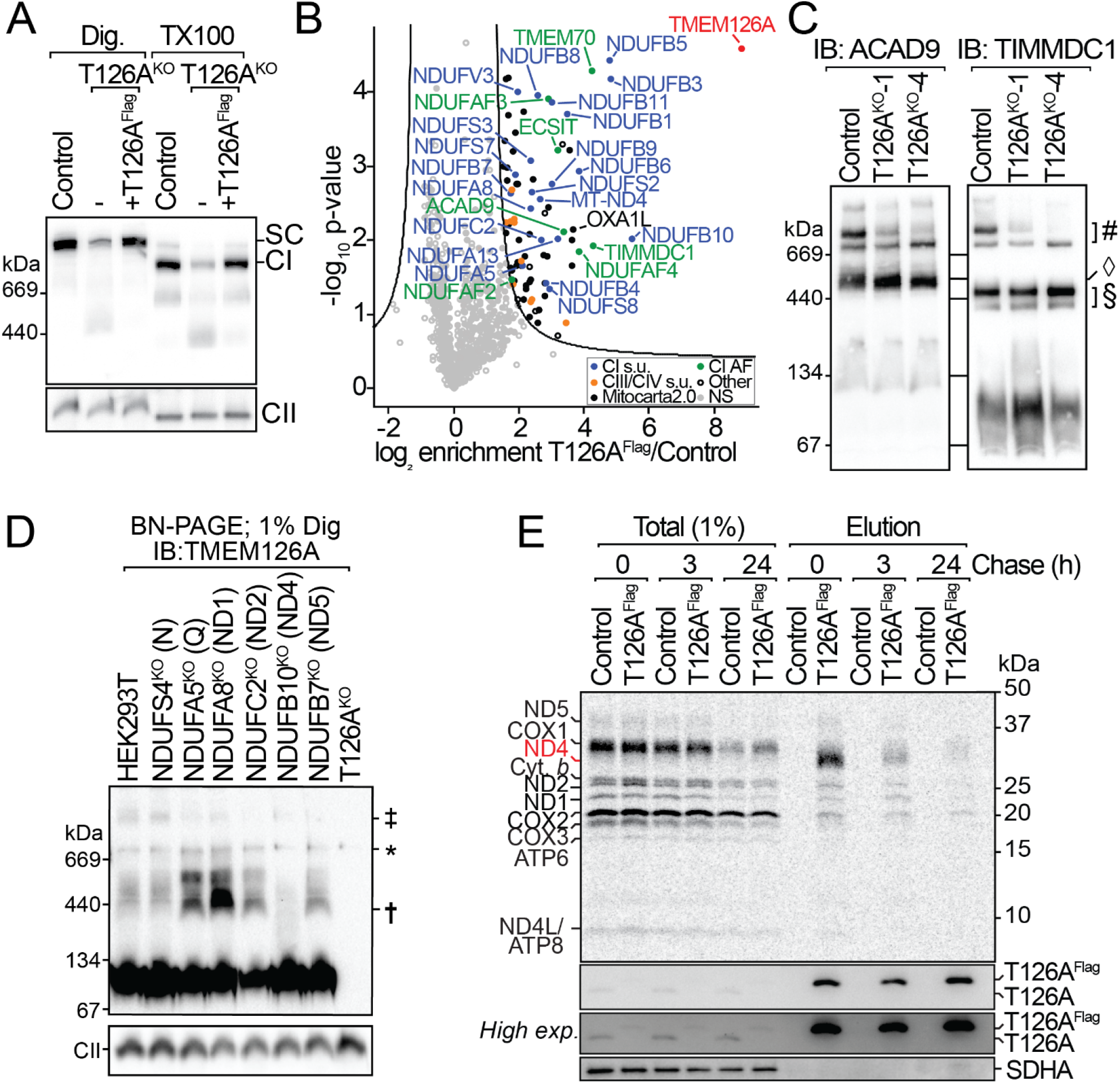
TMEM126A interacts with the ND4-module during complex I assembly. **(A)** Mitochondria isolated from control, T126A^KO^-1 and T126A^KO^-1 +TMEM126A^Flag^ cells and were subjected to BN-PAGE and western blot analysis. Complex II (SDHA) was used as a loading control. **(B)** Digitonin solubilized mitochondria from control and T126A^KO^-1 +TMEM126A^Flag^ expressing cells were subjected to affinity enrichment using anti-Flag agarose beads. Elutions were subjected to LFQ proteomics. Line represents FDR<1%. **(C)** Mitochondria from control and T126A^KO^ cells were solubilized in digitonin and analysed by BN-PAGE and immunoblotting using antibodies as indicated. ◊, ~450 kDa MCIA complex; §, 400/400 kDa TIMMDC1 complex; #, 680/720 kDa complexes **(D)** Mitochondria from control and complex I accessory subunit KO cells were isolated and analysed by BN-PAGE and western blotting using TMEM126A antibodies. A high exposure blot with saturated low molecular weight signal is shown. Complex II (SDHA) was used as a loading control. †, Accumulated TMEM126A complex; ‡, High molecular weight complex; *, non-specific signal. **(E)** Pulse labelled mtDNA-encoded subunits from control and T126A^KO^-1 +TMEM126A^Flag^ expressing cells were chased for the indicated times, Isolated mitochondria were solubilized in digitonin and subjected to affinity enrichment and SDS-PAGE analysis and phosphorimaging. Immunoblotting for TMEM126A and SDHA served as controls.

To investigate TMEM126A-containing complexes in more detail, we investigated changes using representative complex I accessory subunit knockout cell lines of each module (Stroud et al., 2016) (**Fig. 3D**). In control cells, TMEM126A mainly migrated as a low molecular weight species with a portion of it forming higher molecular weight complexes ~440kDa to ~1MDa in size (**Fig. 3D**). These complexes were the same in NDUFS4^KO^ mitochondria, suggesting that they are independent of the N-module. Analysis of NDUFA5^KO^, NDUFA8^KO^, NDUFC2^KO^ and NDUFB7^KO^, which represent the Q-, ND1, ND2 and ND5-modules respectively, showed an accumulation of the ~440kDa complexes and a concomitant reduction in the higher molecular weight complex. This suggests that defects in the assembly of the membrane arm of complex I affect TMEM126A-containing complexes. Moreover, the ~440 kDa complexes were lost in NDUFB10^KO^ mitochondria (representing the ND4-module). Since all other subunits of the ND4-module are lost in NDUFB10^KO^ (Stroud et al., 2016), this suggests that TMEM126A-containing complexes are dependent on subunits of ND4-module for their formation.

Given these results, we assessed whether TMEM126A^Flag^ engages with specific, newly-synthesised mtDNA-encoded subunits We performed pulse-labelling of mtDNA-encoded proteins in cells using [35S]-Methionine in the presence of anisomycin (to block cytosolic translation) for 2 hours, before removal of the radiolabel and anisomycin for up to 24 hours to allow for maturation of the subunits and their assembly into the OXPHOS complexes (Formosa et al., 2016). Flag co-immunoprecipitation and SDS-PAGE analysis revealed a strong but transient enrichment of TMEM126A^Flag^ with the mtDNA-encoded complex I subunit ND4 (**Fig. 3E**). The ND4 subunit was not clearly identified in ‘total’ samples, indicating that the enrichment of this protein by TMEM126A^Flag^ was highly efficient. The relatively minor association between TMEM126A^Flag^ with ND1 and ND2 is also consistent with the proteomic analysis and suggests that TMEM126A is involved in assembly of the ND4-module even during integration with the other assembly modules (Stroud et al., 2016). As expected for an assembly factor, the interaction between TMEM126A and the ND4-module is transient since it is not part of the final assembled complex I. Taken together, these results demonstrate that TMEM126A is an assembly factor for the ND4-module.

### TMEM126A functions independently of TMEM126B in complex I assembly

TMEM126A and its paralog TMEM126B diverged at the root of mammalian evolution; homologs of TMEM126A are present in metazoans while TMEM126B appears confined to mammals (Elurbe and Huynen, 2016). We questioned whether the residual assembly of complex I in TMEM126A deficient cells may be due to the presence of TMEM126B playing a compensatory role. If so, we reasoned that overexpression of TMEM126B in T126A^KO^ cells would enhance complex I assembly. However, additional expression of TMEM126B^Flag^ was not able to rescue complex I levels in T126A^KO^ mitochondria (**Fig. 4A; Fig. S1A**), suggesting that TMEM126A and TMEM126B do not have overlapping functions. Expression of TMEM126A^Flag^ was able complement this assembly defect as expected (**Fig. 4A; Fig. S1B**).

**Figure 4:**
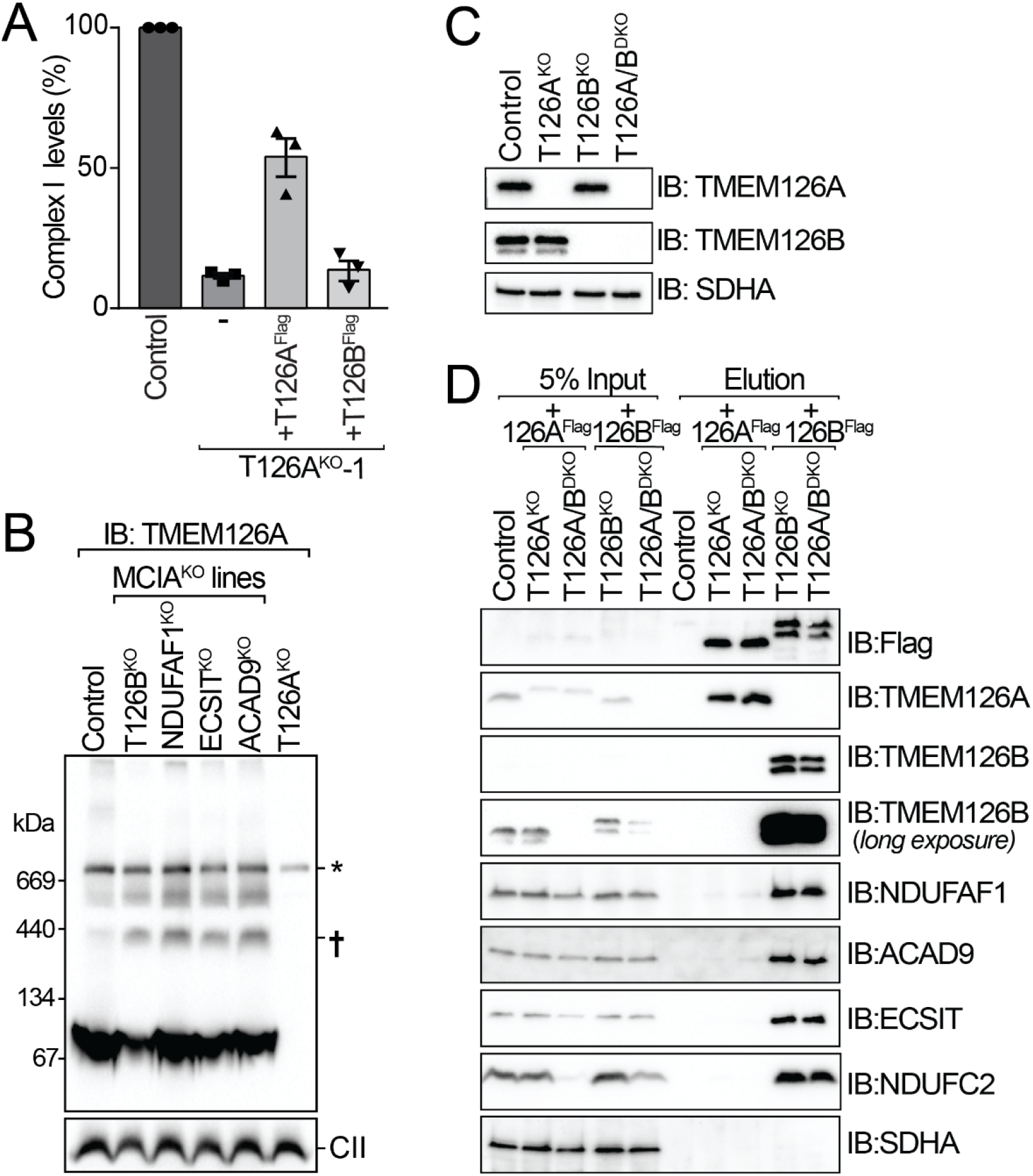
TMEM126A functions independently of TMEM126B. **(A)**The relative abundance of fully assembled complex I was quantified in control and T126A^KO^ cells expressing TMEM126A^Flag^ and TMEM126B^Flag^ as indicated (mean ± SEM, n=3). **(B)**Mitochondria from MCIA^KO^ and T126A^KO^ cells were analysed by BN-PAGE following 1% digitonin solubilization and immunoblotting using TMEM126A antibodies. CII (SDHA) was used as a loading control. †, accumulated TMEM126A complex; *, non-specific signal. **(C)**Mitochondria were isolated from control, T126A^KO^, T126B^KO^ and T126A/B^DKO^ cells and analysed by SDS-PAGE as western blotting with antibodies as shown. SDHA was used as a loading control. **(D)**Mitochondria were isolated from cells expressing TMEM126A^Flag^ or TMEM126B^Flag^ as indicated, solubilized in 1% digitonin and subjected to Flag affinity enrichment prior to SDS-PAGE analysis. Immunoblotting was performed with antibodies as indicated.

While TMEM126B does not complement the loss of TMEM126A, it has been posed that the reverse may be true. Using proteomic analysis of BN-PAGE gel slices from fibroblasts with pathogenic *TMEM126B* mutations, Sánchez-Caballero and colleagues (2016) found that TMEM126A co-migrated with the remaining components of the MCIA complex. This led to the conclusion that TMEM126A plays a compensatory role in the absence of TMEM126B (Sánchez-Caballero et al., 2016b). Indeed, we also observed an accumulation of a TMEM126A complex in TMEM126B knockout mitochondria (**Fig. 4B**; †). However, when we assessed mitochondria lacking MCIA core subunits (NDUFAF1^KO^, ECSIT^KO^ or ACAD9^KO^ lines), which leads to complete loss of the MCIA complex (Formosa et al., 2020), TMEM126A complexes were still present and accumulated even further (**Fig. 4B**). Thus, TMEM126A-containing complexes are independent of the MCIA and the faster migrating TMEM126A complex represents an independent species that accumulates due to a disruption in complex I assembly. Finally, we generated a TMEM126A/TMEM126B double knockout (T126A/B^DKO^) cell line and found that the stabilities of TMEM126A and TMEM126B are independent of each other (**Fig. 4C; Table S1**). We expressed TMEM126A^Flag^ or TMEM126B^Flag^ in the T126A/B^DKO^ cell line and performed co-immunoprecipitation and SDS-PAGE analysis (**Fig. 4D**). TMEM126B^Flag^, but not TMEM126A^Flag^ efficiently enriched MCIA subunits ACAD9, ECSIT and NDUFAF1 along with the complex I subunit NDUFC2. This indicates that in the absence of TMEM126B, TMEM126A does not further engage the MCIA complex. We conclude that TMEM126A and TMEM126B do not play compensatory roles in the absence of the other protein, but rather, are involved directly in the assembly and stability of different stages in complex I assembly. TMEM126A is therefore a newly identified assembly factor for the ND4-module of complex I and functions independently of its paralog TMEM126B.

## Discussion

### TMEM126A is important for biogenesis of mitochondrial complex I through ND4-module assembly

The analysis of proteins involved in mitochondrial disease where no clear molecular function is ascribed is critical to improving diagnosis and, potentially, future patient outcomes. Mutations in *TMEM126A* have been identified in numerous cases of autosomal recessive optic atrophy (Desir et al., 2012; Hanein et al., 2009; Kloth et al., 2019; La Morgia et al., 2019), however a detailed molecular function of this protein has been lacking. From the reported cases of patients with mutations in TMEM126A that result in optic atrophy, one patient was reported to have a ‘partial deficiency of complex I’ in patient-derived fibroblasts (Hanein et al., 2009). While an MRI conducted was considered normal, the patient was reported to have moderate hypertrophic cardiomyopathy, which is a more common presentation for patients with complex I deficiency (Frazier et al., 2020; Frazier et al., 2019; Gorman et al., 2016; Hanein et al., 2009). Analysis of TMEM126A-interacting proteins revealed an association with TMEM70, a protein long known to play a role in assembly of ATP synthase through incorporation of subunit c into the complex V rotor (Kovalčíková et al., 2019; Spiegel et al., 2011), but more recently also implicated in assembly of the complex I ND4-module (PD-a intermediate) (Guerrero-Castillo et al., 2017; Sánchez-Caballero et al., 2020).

Assembly factors have been found to play important roles during the biogenesis of complex I, including the delivery of co-factors (Sheftel et al., 2009), post-translational modification of subunits (Rhein et al., 2013; Rhein et al., 2016; Zurita Rendon et al., 2014), the insertion of subunits into the inner membrane and stabilization of incompletely assembled modules (Formosa et al., 2018). Previous studies have started to elucidate a clearer picture for the assembly of the distal membrane arm of complex I, with a number of assembly factors identified to participate in the biogenesis of this region including FOXRED1, TMEM70, DMAC1 and DMAC2 (Formosa et al., 2015; Guerrero-Castillo et al., 2017; Stroud et al., 2016). Indeed, complexome profiling has previously demonstrated that DMAC2, FOXRED1 and TMEM70 co-migrate with subunits of the ND4-module of complex I during assembly (Guerrero-Castillo et al., 2017). Surprisingly, TMEM126A was not detected with these complex I assembly intermediates (Guerrero-Castillo et al., 2017). Interestingly, of these ND4-module assembly factors, TMEM126A only enriched TMEM70 and not FOXRED1 or DMAC2, suggesting that while they participate in assembly of the same module, they may not participate during the same stage of this process. Indeed, our analysis demonstrated a strong enrichment of newly-synthesised ND4 protein associated with TMEM126A. Also, TMEM126A-containing complexes were strongly disrupted in NDUFB10^KO^ mitochondria - a subunit belonging to the ND4-module whose loss leads to turnover of all ND4-module subunits including ND4 itself (Fiedorczuk et al., 2016; Stroud et al., 2016). The clinical link between TMEM126A and assembly of the ND4-module is further supported by the numerous patients with the maternally inherited mtDNA m.11778G>A mutation in the *MT-ND4* gene, the most common mtDNA mutation that leads to Leber’s Hereditary Optic Neuropathy (LHON), which is a similar degenerative condition that affects the optic nerve neurons (Brown et al., 2001; Catarino et al., 2017; Newman, 2005; Wallace et al., 1988). We propose that TMEM126A plays a prominent role in the assembly of complex I through association with the ND4-module, and loss of this protein results in decreased levels of complex I leading to optic atrophy. Future genetic diagnosis of patients with complex I deficiency should consider the possibility of genetic variants in the *TMEM126A* gene as a possible cause of mitochondrial disease.

### Duplication and specialization of the TMEM126 locus

The ancestral *TMEM126A* gene appeared to undergo a duplication event giving rise to TMEM126A and TMEM126B, both having a role in the assembly of complex I (Elurbe and Huynen, 2016). Given the recent duplication at the root of mammalian evolution, these genes have undergone rapid adaptation and associate with distinct membrane assembly intermediates of complex I - TMEM126B as part of the MCIA complex required for ND2-module (Formosa et al., 2020) and here, we show that TMEM126A associates with ND4. Both the ND2 and ND4 subunits are membrane proteins that have evolved from the ancient Na+/H+ Mrp antiporter complex (Sazanov, 2015). This alludes to some interesting questions for non-mammalian metazoans where only TMEM126A is found. Analysis of complex I assembly defects in fly muscle upon knockdown of the subunits *Drosophila* (d)NDUFV1 or dNDUFS5 revealed the presence of a complex I intermediate consisting of the distal portion of the membrane arm of complex I (including the ND4-module), conserved components of the MCIA complex as well as the *Drosophila* homologue of TMEM126A, termed dTMEM126B (Garcia et al., 2017). Might this ancestral protein be involved in assembly of two distinct complex I modules? Interestingly, in addition to TMEM126A and TMEM126B sharing an evolutionary origin, the transmembrane proteins TMEM70 (involved in ND4-module assembly) and TMEM186 (involved in ND2-module assembly by interacting with ND3) also share a common ancestor (Formosa et al., 2020; Guerrero-Castillo et al., 2017; Sánchez-Caballero et al., 2020). This may suggest potential symmetry involved in the assembly and integration of the ND2- and ND4-modules. From an evolutionary perspective, further studies could provide useful insights into how complex I assembly has adapted over time. It is also unclear why loss of TMEM126A from mammalian cells appears to be less severe than the loss of TMEM126B, given that TMEM126A came first. Indeed, individuals with mutations in TMEM126A primarily present with optic atrophy and most have been identified with homozygous nonsense mutations (Hanein et al., 2009; Meyer et al., 2010). This is in contrast to individuals with TMEM126B mutations who present with adult onset myopathy and all identified cases have at least one missense mutation, suggesting total ablation of TMEM126B may not be compatible with human life (Alston et al., 2016; Sánchez-Caballero et al., 2016b). One possibility is that TMEM126B plays a crucial role in the MCIA complex for ND2-module assembly, while other (unidentified) mitochondrial machineries may be able to compensate for the loss of TMEM126A for ND4-module assembly. Further studies into the role of TMEM126A as a new component of the complex I assembly machinery and its evolutionary relationship with TMEM126B are warranted to uncover the mechanism behind these observations.

## Acknowledgements

We thank the Bio21 Mass Spectrometry and Proteomics Facility (MMSPF) and the Monash Proteomics and Metabolomics Facility (MPMF) for the provision of instrumentation, training, and technical support and Monash Flowcore for cell sorting. We acknowledge funding from the National Health and Medical Research Council (NHMRC Project Grants 1164459 and 1165217 to MTR; 1125390 to MTR and DAS; NHMRC Fellowship 1140851 to DAS).

## Methods and Materials

### Cell Culturing

HEK293T cells (ATCC; CRL-3216) were cultured in Dulbecco’s Modified Eagle’s Medium (DMEM) supplemented with 10% (v/v) Fetal Bovine Serum (FBS; Life Technologies; 10499-044), 1% penicillin-streptomycin (P/S; Sigma-Aldrich; P3358), 1× GlutaMAX^™^ (Life Technologies; 35050061) and 50 μg/mL uridine (Sigma-Aldrich; U6381). For galactose screening of the TMEM126A/126B^DKO^ cell line, glucose-free DMEM (Life Technologies; 11966-025) was supplemented with 10% (v/v) FBS, 25 mM d-(+)-galactose (Sigma-Aldrich; G0750), 1% P/S, 1× GlutaMAX^™^, 1 mM sodium pyruvate (Life Technologies; 11360070), and 50 μg/mL uridine. All cells were incubated at 37°C with 5% CO_2_ in a humidified environment.

For SILAC experiments, cells were cultured as previously described (Stroud et al., 2016). Knockout cell lines and their respective controls were grown in DMEM without lysine and arginine, supplemented with 10% (v/v) dialyzed FBS (GE Healthcare; SH30079.03), 1% P/S, 1× GlutaMAX^™^ Supplement, 1 mM sodium pyruvate, and 50 μg/mL uridine (Sigma-Aldrich; U6381). Media also contained either ‘light’ lysine and arginine (Lysine; Sigma-Aldrich; L5626, Arginine; Sigma-Aldrich; A5131) or ‘heavy’ ^13^C_61_^5^N_2_-Lysine (Silantes; 211604102) and ^13^C_6_^15^N_4_-Arginine (Cambridge Isotope Labs Inc; CNLM-539-H-1). Cells were cultured in SILAC ‘light’ or ‘heavy’ media for two weeks prior to harvesting by centrifugation (800*g*, 5 min, 4°C), followed by mitochondrial isolation and quantitative proteomics (described below).

### Cloning and Molecular Biology

CRISPR/Cas9 gene editing was performed using the pSpCas9(BB)-2A-GFP (X458) plasmid construct (a gift from F. Zhang; Addgene; plasmid 48138 (Ran et al., 2013)). Gene-specific target sites and guide RNAs were designed using the web-based tool, CHOPCHOP (Montague et al., 2014). Targeting strategies and gRNA sequences are detailed in **Table S1**. For the generation of rescue cell lines, cDNA from target genes were amplified from a HEK293T cDNA library with a 3’ Flag sequence incorporated into the reverse primer. The constitutively expressed retroviral vector pBABE-puro (Addgene; plasmid 1764; (Morgenstern and Land, 1990)) was used to express the cDNA of interest. Sequence verification of all plasmid constructs was performed by Micromon Genomics (Monash University).

### Transfection and Stable Cell Line Generation

Cells were transfected using Lipofectamine^™^ LTX and PLUS^™^ Reagents (Life Technologies; 15338100) according to the manufacturer’s instructions. Stable cell lines expressing the protein of interest were generated by transfecting HEK293T cells with pBABE-puro constructs encoding the desired cDNA. Retroviral supernatants were harvested after 48 hours, filtered through a 0.45 μm filter (Merck; SLHV033RS) and used for transduction of knockout cell lines with the addition of 8 μg/mL polybrene. Infected cells were selected for by the addition of 2 μg/mL puromycin (InvivoGen; ant-pr). Expression was verified by SDS-PAGE and/or BN-PAGE analysis.

### Gene Editing and Screening

Clonal knockout cell populations were generated by fluorescence-activated cell sorting (FACS) for GFP expression (encoded in the CRISPR/Cas9 construct) 24 hours post transfection. FACS analysis and single-cell sorting was performed by Flowcore (Monash University). Loss of TMEM126A was screened by PCR of extracted gDNA and subsequent western blotting analysis. Respiratory complex I deficient TMEM126A/B^DKO^ clones were identified by a growth defect on galactose-based DMEM and subsequent western blotting analysis. Knockout cell lines were verified by sequence identification of individual indels (**Table S1**) as previously described (Stroud et al., 2013). Complete loss of a protein product was further validated by SDS-PAGE and/or BN-PAGE. Other complex I accessory subunit and assembly factor KO lines are described elsewhere (**Table S1**)(Formosa et al., 2020; Stroud et al., 2016).

### Mitochondrial Isolation, Gel Electrophoresis and Immunoblotting

Mitochondria were isolated from cultured cells by differential centrifugation as previously described (Formosa et al., 2020; Johnston et al., 2002) Quantitation of protein concentration was estimated by bicinchoninic acid (BCA) assay using the Thermo Scientific^™^ Pierce^™^ BCA Protein Assay kit (Thermo Fisher Scientific; 23227).

Polyacrylamide gel electrophoresis techniques were performed as previously described (Formosa et al., 2020; McKenzie et al., 2007; Schagger and von Jagow, 1987; Wittig et al., 2006). For SDS-PAGE, crude mitochondrial samples were solubilised in 2× SDS loading dye (100 mM Tris-Cl pH 6.8, 200 mM DTT, 4% (w/v) SDS, 20% (v/v) glycerol, 0.1% bromophenol blue) or 1× LDS Sample Buffer (Thermo Fisher Scientific; NP0008) supplemented with 100 mM DTT. For western blotting, 10% acrylamide gels were used while continuous 10-16.5% gels were used for separation of radiolabeled proteins. For BN-PAGE analysis of mitochondrial complexes, crude mitochondria were solubilised in 1% digitonin or 1% Triton X-100 in solubilisation buffer (20 mM Bis-Tris pH 7.0, 50 mM NaCl, 10% (v/v) glycerol) for 10 min on ice, followed by centrifugation at 16,000 *g* for 10 min. The supernatant was separated from the pellet of unsolubilised material and mixed with 10× BN loading dye (500 mM ∊-amino n-caproic acid, 100 mM Bis-Tris pH 7.0, 5% (w/v) Coomassie Brilliant Blue G-250) to a final 1× concentration. Western transfer onto Immoiblon-P PVDF membranes (Merck Millipore; IPVH00010) was performed using the Power Blotter XL System (Thermo Fisher Scientific; PB0013) or Novex™ Semi-Dry Blotter (Thermo Fisher Scientific; SD1000) in accordance with the manufacturer’s instructions. Blots were stained in PVDF stain (50% (v/v) methanol, 7% (v/v) acetic acid, 0.1% (w/v) Coomassie Brilliant Blue R-250) and excess Coomassie was removed by incubating in PVDF destain (50% (v/v) methanol, 7% (v/v) acetic acid). PVDF stain was completely removed with 100% methanol and blots were subsequently washed in Tris-buffered Saline (TBS; 20 mM Tris-Cl pH 7.5, 150 mM NaCl) with 0.1% Tween-20 (TBS-T). The primary antibodies used in this study are as follows: ACAD9 (in-house; (Formosa et al., 2020)), ATP5A (Abcam; ab14748), Core 1 (Thermo Fisher Scientific; 459140), COX4 (Abcam; ab110261), ECSIT (in-house; (Formosa et al., 2020)), Flag (M2 clone; Sigma-Aldrich; F1804), NDUFA9 (in-house; (Stroud et al., 2013)), NDUFAF1 (in-house; (Formosa et al., 2020)), NDUFB8 (Abcam; ab110242), NDUFB11 (Abcam; ab183716), NDUFC2 (Santa Cruz Biotechnology; sc-398719), NDUFS5 (in-house), SDHA (Abcam; ab14715), TIMMDC1 (Sigma-Aldrich; HPA053214), TMEM126A (Sigma-Aldrich; HPA046648), TMEM126B (Sigma-Aldrich; HPA014480). Horseradish peroxidase coupled secondary antibodies (anti-mouse IgG; Sigma-Aldrich; A9044, anti-rabbit-IgG; Sigma-Aldrich; A0545) were used with Clarity™ Western ECL Substrate (Bio-Rad; 1705061) and Clarity Max™ Western ECL Substrate (Bio-Rad; 1705062) for detection with the ChemiDocTM XRS+ System (Bio-Rad).

For densitometric analysis, western blot exposures were analysed in ImageLab software (Bio-Rad) by drawing a box around the region of interest as well as a separate region for background correction. The same sized box was the used for all samples to be analysed. Pixel intensity was also measured for a loading control. The normalized signal intensity was then taken as a percentage of control samples and analysed using GraphPad Prism (v7.01).

### Radiolabeling of mtDNA-encoded proteins and Affinity Enrichment analysis

Cells were cultured for radiolabeling as previously described (Formosa et al., 2016; Formosa et al., 2020). To summarise, cells were labelled in Met/Cys-free DMEM (Life Technologies; 21013024), supplemented with 10% (v/v) dialysed FBS, 1% P/S, 1× GlutaMAX^™^, 1 mM sodium pyruvate, 50 μg/mL uridine, 7 μg/mL anisomycin (Sigma-Aldrich; A9789), and 7 μCi [^35^S]-Methionine/cysteine (PerkinElmer; NEG072007MC) and labelled for 2 hours. Labelling was quenched with 10 μM ‘cold’ methionine (Sigma-Aldrich; M9625) and replaced with standard DMEM growth media for 0, 3 or 24 hours. Following this, cells were washed with PBS and media was replaced with regular DMEM. Cells were then chased for 0, 3 and 24 hours. Cells were harvested by centrifugation (800*g*, 5 min, 4°C). Pellets were stored at −20°C until use. Isolated mitochondria from radiolabelled cells were used for Flag affinity enrichment and SDS-PAGE analysis and immunoblotting. Detection of radiolabelled mt-DNA subunits was performed using an Amersham^™^ Typhoon^™^ Biomolecular Imager (GE Healthcare).

For affinity enrichment of Flag-tagged proteins, mitochondria were solubilised with 1% digitonin in solubilisation buffer (20 mM Bis-Tris pH 7.0, 50 mM NaCl, 10% (v/v) glycerol) for 10 min on ice. Solubilised mitochondria were then clarified by centrifugation at 16,000 *g* for 10 min and incubated with pre-washed anti-Flag^®^ M2 Affinity Gel (Merck; A2220) at 4 °C for 2 hours. Following affinity purification, Flag affinity gel was washed with 0.1% digitonin in solubilisation buffer. For samples used for proteomics or western blotting analysis, Flag-tagged proteins were eluted by incubation with 150 μg/mL Flag tag peptide (Merck; F3290) with 0.1% digitonin in solubilisation buffer for 30 min at 4°C. Eluates were then prepared for SDS-PAGE by the addition of 4 × SDS loading dye or for proteomics analysis as described below. For pulse-chase experiments, enriched proteins were removed from the Flag affinity gel by incubating beads in 2× SDS loading dye (100 mM Tris-Cl pH 6.8, 200 mM DTT, 4% (w/v) SDS, 20% (v/v) glycerol, 0.1% bromophenol blue) at room temperature for 15 min and separated from the affinity gel by centrifugation at 16,000 *g* for 10 min.

### Proteomics sample preparation

For SILAC analysis, control and T126A^KO^ cells were cultured in ‘Heavy’ and ‘Light’ SILAC media as described above. Sample preparation was performed as previously described with some modifications (Dibley et al., 2020; Formosa et al., 2020; Stroud et al., 2016). Briefly, protein concentrations were determined by BCA and equal amounts of differently labeled control and T126A^KO^ cells were mixed and mitochondria isolated as described. Two samples were heavy labeled T126A^KO^ cells and light control labeled cells, while a third consisted of a label switch. Following mitochondrial isolation, samples were solubilized in 1% sodium deoxycholate, 100 mM Tris-Cl pH 8.1, 40 mM chloroacetamide and 10 mM TCEP prior to vortexing and heating for 5 minutes at 99 °C with vigorous shaking at 1400 rpm. Samples were then sonicated in a water bath at room temperature for 15 minutes. Proteomic digestion was performed by the addition of 1 μg proteomics-grade Trypsin (Promega) and incubated overnight at 37 °C. Supernatants were mixed with Ethyl acetate (99%) and TFA (1%), then transferred to 3x 14G 3M^™^ Empore^™^ SDB-RPS stage tips (Kulak et al., 2014). Samples were then centrifuged through the stage tip at 3000 *g* at room temperature (Kulak et al., 2014). Stage tips were then washed with 99% ethyl acetate with 1% TFA, followed by a second wash with 99% ethyl acetate and 0.1% TFA. Samples were eluted with 80% ACN and 1% NH_4_OH and acidified to a final TFA concertation 1% before drying samples down in a SpeedVac.

For affinity enrichment sample preparation, elutions were precipitated using five volumes of ice-cold acetone and incubated at −20 °C overnight. Samples were centrifuged at 16,000 × g for 10 min at 4 °C, following which the precipitated pellet was solubilized in 8 M urea and 50 mM ammonium bicarbonate. This was sonicated in a water bath at room temperature for 15 min before addition of 5 mM Tris(2-carboxyethyl)phosphine hydrochloride (TCEP) and 50 mM chloroacetamide (Sigma-Aldrich) and incubated at 37 °C while shaking. The sample was then treated with ammonium bicarbonate to dilute the urea to a final concentration of 2 M. Sequencing grade modified trypsin (1 μg; Promega; V5113) was added to the sample before incubation overnight at 37 °C. After this the sample was acidified with 10% trifluoroacetic acid (TFA) to a final concentration of 1%. Stage tips were generated with two plugs of 3 M^™^ Empore^™^ SDB-XC Extraction Disks (Fisher Scientific) that were activated with 100% acetonitrile (ACN) via centrifugation. All spins were performed at 1800 × g. The tips were washed with 0.1% TFA, 2% ACN three times. The sample was added to the stage tip and eluted with 80% ACN and 0.1% TFA. The eluates were subsequently dried down using a SpeedVac.

### Proteomics analysis

Peptides were reconstituted in 2% ACN, 0.1% TFA and transferred to autosampler vials for analysis by online nano-HPLC/electrospray ionization-MS/MS using an Orbitrap Fusion Lumos instrument connected to an Ultimate 3000 HPLC using the instrument settings described in detail in (Formosa et al., 2020). Raw files were analysed using the MaxQuant platform (Tyanova et al., 2016a) version 1.6.5.0 searching against the UniProt human database containing reviewed, canonical entries (January 2019) and a database containing common contaminants. Parameters used in the SILAC and LFQ searches are described in Formosa et al. (2020). Statistical analysis was performed using the Perseus platform (Tyanova et al., 2016b) version 1.6.7.0 as described in Formosa et al. (2020). Log_2_-transformed median SILAC ratios were mapped on homologous subunits of the respiratory chain complexes. The mass spectrometry proteomics data will be deposited in the ProteomeXchange Consortium via the PRIDE (Perez-Riverol et al., 2019) partner repository at the time of publication.

## References

Alston, C.L., Compton, A.G., Formosa, L.E., Strecker, V., Oláhová, M., Haack, T.B., Smet, J., Stouffs, K., Diakumis, P., Ciara, E., et al. (2016). Biallelic Mutations in TMEM126B Cause Severe Complex I Deficiency with a Variable Clinical Phenotype. American Journal of Human Genetics 99, 217–227.

Andrews, B., Carroll, J., Ding, S., Fearnley, I.M., and Walker, J.E. (2013). Assembly factors for the membrane arm of human complex I. Proceedings of the National Academy of Sciences of the United States of America 110, 18934–18939.

Brown, M.D., Allen, J.C., Van Stavern, G.P., Newman, N.J., and Wallace, D.C. (2001). Clinical, genetic, and biochemical characterization of a Leber hereditary optic neuropathy family containing both the 11778 and 14484 primary mutations. American Journal of Medical Genetics 104, 331–338.

Carelli, V., La Morgia, C., Valentino, M.L., Barboni, P., Ross-Cisneros, F.N., and Sadun, A.A. (2009). Retinal ganglion cell neurodegeneration in mitochondrial inherited disorders. Biochim Biophys Acta 1787, 518–528.

Catarino, C.B., Ahting, U., Gusic, M., Iuso, A., Repp, B., Peters, K., Biskup, S., von Livonius, B., Prokisch, H., and Klopstock, T. (2017). Characterization of a Leber’s hereditary optic neuropathy (LHON) family harboring two primary LHON mutations m.11778G>A and m.14484T>C of the mitochondrial DNA. Mitochondrion 36, 15–20.

De Vries, D.D., Went, L.N., Bruyn, G.W., Scholte, H.R., Hofstra, R.M., Bolhuis, P.A., and van Oost, B.A. (1996). Genetic and biochemical impairment of mitochondrial complex I activity in a family with Leber hereditary optic neuropathy and hereditary spastic dystonia. Am J Hum Genet 58, 703–711.

Desir, J., Coppieters, F., Van Regemorter, N., De Baere, E., Abramowicz, M., and Cordonnier, M. (2012). TMEM126A mutation in a Moroccan family with autosomal recessive optic atrophy. Molecular vision 18, 1849–1857.

Dibley, M.G., Formosa, L.E., Lyu, B., Reljic, B., McGann, D., Muellner-Wong, L., Kraus, F., Sharpe, A.J., Stroud, D.A., and Ryan, M.T. (2020). The Mitochondrial Acyl-carrier Protein Interaction Network Highlights Important Roles for LYRM Family Members in Complex I and Mitoribosome Assembly. Molecular & cellular proteomics: MCP 19, 65–77.

Elurbe, D.M., and Huynen, M.A. (2016). The origin of the supernumerary subunits and assembly factors of complex I: A treasure trove of pathway evolution. Biochimica et Biophysica Acta (BBA) - Bioenergetics 1857, 971–979.

Fiedorczuk, K., Letts, J.A., Degliesposti, G., Kaszuba, K., Skehel, M., and Sazanov, L.A. (2016). Atomic structure of the entire mammalian mitochondrial complex I. Nature 538, 406–410.

Formosa, L.E., Dibley, M.G., Stroud, D.A., and Ryan, M.T. (2018). Building a complex complex: Assembly of mitochondrial respiratory chain complex I. Seminars in Cell & Developmental Biology 76, 154–162.

Formosa, L.E., Hofer, A., Tischner, C., Wenz, T., and Ryan, M.T. (2016). Translation and Assembly of Radiolabeled Mitochondrial DNA-Encoded Protein Subunits from Cultured Cells and Isolated Mitochondria. Methods in Molecular Biology (Clifton, N.J.) 1351, 115–129.

Formosa, L.E., Mimaki, M., Frazier, A.E., McKenzie, M., Stait, T.L., Thorburn, D.R., Stroud, D.A., and Ryan, M.T. (2015). Characterization of mitochondrial FOXRED1 in the assembly of respiratory chain complex I. Human Molecular Genetics 24, 2952–2965.

Formosa, L.E., Muellner-Wong, L., Reljic, B., Sharpe, A.J., Jackson, T.D., Beilharz, T.H., Stojanovski, D., Lazarou, M., Stroud, D.A., and Ryan, M.T. (2020). Dissecting the Roles of Mitochondrial Complex I Intermediate Assembly Complex Factors in the Biogenesis of Complex I. Cell reports 31, 107541.

Frazier, A.E., Compton, A.G., Kishita, Y., Hock, D.H., Welch, A.E., Amarasekera, S.S.C., Rius, R., Formosa, L.E., Imai-Okazaki, A., Francis, D., et al. (2020). Fatal Perinatal Mitochondrial Cardiac Failure Caused by Recurrent De Novo Duplications in the ATAD3 Locus. Med.

Frazier, A.E., Thorburn, D.R., and Compton, A.G. (2019). Mitochondrial energy generation disorders: genes, mechanisms, and clues to pathology. Journal of Biological Chemistry 294, 5386–5395.

Garcia, C.J., Khajeh, J., Coulanges, E., Chen, E.I., and Owusu-Ansah, E. (2017). Regulation of Mitochondrial Complex I Biogenesis in Drosophila Flight Muscles. Cell reports 20, 264–278.

Gorman, G.S., Chinnery, P.F., DiMauro, S., Hirano, M., Koga, Y., McFarland, R., Suomalainen, A., Thorburn, D.R., Zeviani, M., and Turnbull, D.M. (2016). Mitochondrial diseases. Nat Rev Dis Primers 2, 16080.

Gu, J., Wu, M., Guo, R., Yan, K., Lei, J., Gao, N., and Yang, M. (2016). The architecture of the mammalian respirasome. Nature 537, 639–643.

Guerrero-Castillo, S., Baertling, F., Kownatzki, D., Wessels, H.J., Arnold, S., Brandt, U., and Nijtmans, L. (2017). The Assembly Pathway of Mitochondrial Respiratory Chain Complex I. Cell Metabolism 25, 1–12.

Hanein, S., Garcia, M., Fares-Taie, L., Serre, V., De Keyzer, Y., Delaveau, T., Perrault, I., Delphin, N., Gerber, S., Schmitt, A., et al. (2013). TMEM126A is a mitochondrial located mRNA (MLR) protein of the mitochondrial inner membrane. Biochimica et Biophysica Acta (BBA) - General Subjects 1830, 3719–3733.

Hanein, S., Perrault, I., Roche, O., Gerber, S., Khadom, N., Rio, M., Boddaert, N., Jean-Pierre, M., Brahimi, N., Serre, V., et al. (2009). TMEM126A, encoding a mitochondrial protein, is mutated in autosomal-recessive nonsyndromic optic atrophy. Am J Hum Genet 84, 493–498.

Heide, H., Bleier, L., Steger, M., Ackermann, J., Dröse, S., Schwamb, B., Zörnig, M., Reichert, A.S., Koch, I., Wittig, I., et al. (2012). Complexome profiling identifies TMEM126B as a component of the mitochondrial complex I assembly complex. Cell Metabolism 16, 538–549.

Johnston, A.J., Hoogenraad, J., Dougan, D.A., Truscott, K.N., Yano, M., Mori, M., Hoogenraad, N.J., and Ryan, M.T. (2002). Insertion and assembly of human tom7 into the preprotein translocase complex of the outer mitochondrial membrane. Journal of Biological Chemistry 277, 42197–42204.

Kloth, K., Synofzik, M., Kernstock, C., Schimpf-Linzenbold, S., Schuettauf, F., Neu, A., Wissinger, B., and Weisschuh, N. (2019). Novel likely pathogenic variants in TMEM126A identified in non-syndromic autosomal recessive optic atrophy: two case reports. BMC Medical Genetics 20, 62.

Kovalčíková, J., Vrbacký, M., Pecina, P., Tauchmannová, K., Nůsková, H., Kaplanová, V., Brázdová, A., Alán, L., Eliáš, J., Čunátová, K., et al. (2019). TMEM70 facilitates biogenesis of mammalian ATP synthase by promoting subunit c incorporation into the rotor structure of the enzyme. FASEB journal: official publication of the Federation of American Societies for Experimental Biology 33, 14103–14117.

Kulak, N.A., Pichler, G., Paron, I., Nagaraj, N., and Mann, M. (2014). Minimal, encapsulated proteomic-sample processing applied to copy-number estimation in eukaryotic cells. Nature Methods 11, 319–324.

La Morgia, C., Caporali, L., Tagliavini, F., Palombo, F., Carbonelli, M., Liguori, R., Barboni, P., and Carelli, V. (2019). First TMEM126A missense mutation in an Italian proband with optic atrophy and deafness. Neurology. Genetics 5, e329.

Lazarou, M., McKenzie, M., Ohtake, A., Thorburn, D.R., and Ryan, M.T. (2007). Analysis of the assembly profiles for mitochondrial-and nuclear-DNA-encoded subunits into complex I. Molecular and Cellular Biology 27, 4228–4237.

Lin, Y.H., Wang, N.K., Yeung, L., Lai, C.C., and Chuang, L.H. (2018). Juvenile open-angle Glaucoma associated with Leber's hereditary optic neuropathy: a case report and literature review. BMC ophthalmology 18, 323.

McKenzie, M., Lazarou, M., Thorburn, D.R., and Ryan, M.T. (2007). Analysis of mitochondrial subunit assembly into respiratory chain complexes using Blue Native polyacrylamide gel electrophoresis. Analytical biochemistry 364, 128–137.

Meyer, E., Michaelides, M., Tee, L.J., Robson, A.G., Rahman, F., Pasha, S., Luxon, L.M., Moore, A.T., and Maher, E.R. (2010). Nonsense mutation in TMEM126A causing autosomal recessive optic atrophy and auditory neuropathy. Molecular vision 16, 650–664.

Montague, T.G., Cruz, J.M., Gagnon, J.A., Church, G.M., and Valen, E. (2014). CHOPCHOP: a CRISPR/Cas9 and TALEN web tool for genome editing. Nucleic Acids Research 42, W401–407.

Morgenstern, J.P., and Land, H. (1990). Advanced mammalian gene transfer: high titre retroviral vectors with multiple drug selection markers and a complementary helper-free packaging cell line. Nucleic Acids Research 18, 3587–3596.

Newman, N.J. (2005). Hereditary optic neuropathies: from the mitochondria to the optic nerve. American journal of ophthalmology 140, 517–523.

Perez-Riverol, Y., Csordas, A., Bai, J., Bernal-Llinares, M., Hewapathirana, S., Kundu, D.J., Inuganti, A., Griss, J., Mayer, G., Eisenacher, M., et al. (2019). The PRIDE database and related tools and resources in 2019: improving support for quantification data. Nucleic Acids Res 47, D442–d450.

Ran, F.A., Hsu, P.D., Wright, J., Agarwala, V., Scott, D.A., and Zhang, F. (2013). Genome engineering using the CRISPR-Cas9 system. Nature Protocols 8, 2281–2308.

Rhein, V.F., Carroll, J., Ding, S., Fearnley, I.M., and Walker, J.E. (2013). NDUFAF7 methylates arginine 85 in the NDUFS2 subunit of human complex I. Journal of Biological Chemistry 288, 33016–33026.

Rhein, V.F., Carroll, J., Ding, S., Fearnley, I.M., and Walker, J.E. (2016). NDUFAF5 Hydroxylates NDUFS7 at an Early Stage in the Assembly of Human Complex I. Journal of Biological Chemistry 291, 14851–14860.

Sánchez-Caballero, L., Elurbe, D.M., Baertling, F., Guerrero-Castillo, S., van den Brand, M., van Strien, J., van Dam, T.J.P., Rodenburg, R., Brandt, U., Huynen, M.A., et al. (2020). TMEM70 functions in the assembly of complexes I and V. Biochimica et Biophysica Acta (BBA) - Bioenergetics 1861, 148202.

Sánchez-Caballero, L., Guerrero-Castillo, S., and Nijtmans, L. (2016a). Unraveling the complexity of mitochondrial complex I assembly: A dynamic process. Biochimica et biophysica acta 1857, 980–990.

Sánchez-Caballero, L., Ruzzenente, B., Bianchi, L., Assouline, Z., Barcia, G., Metodiev, M.D., Rio, M., Funalot, B., van den Brand, M.A.A., Guerrero-Castillo, S., et al. (2016b). Mutations in Complex I Assembly Factor TMEM126B Result in Muscle Weakness and Isolated Complex I Deficiency. American Journal of Human Genetics 99, 208–216.

Sazanov, L.A. (2015). A giant molecular proton pump: structure and mechanism of respiratory complex I. Nature reviews. Molecular cell biology 16, 375–388.

Schagger, H., and von Jagow, G. (1987). Tricine-sodium dodecyl sulfate-polyacrylamide gel electrophoresis for the separation of proteins in the range from 1 to 100 kDa. Analytical biochemistry 166, 368–379.

Schlattner, U., Tokarska-Schlattner, M., and Wallimann, T. (2006). Mitochondrial creatine kinase in human health and disease. Biochimica et biophysica acta 1762, 164–180.

Sheftel, A.D., Stehling, O., Pierik, A.J., Netz, D.J., Kerscher, S., Elsasser, H.P., Wittig, I., Balk, J., Brandt, U., and Lill, R. (2009). Human ind1, an iron-sulfur cluster assembly factor for respiratory complex I. Molecular and Cellular Biology 29, 6059–6073.

Signes, A., and Fernandez-Vizarra, E. (2018). Assembly of mammalian oxidative phosphorylation complexes I-V and supercomplexes. Essays in biochemistry 62, 255–270.

Spiegel, R., Khayat, M., Shalev, S.A., Horovitz, Y., Mandel, H., Hershkovitz, E., Barghuti, F., Shaag, A., Saada, A., Korman, S.H., et al. (2011). TMEM70 mutations are a common cause of nuclear encoded ATP synthase assembly defect: further delineation of a new syndrome. Journal of medical genetics 48, 177–182.

Spinelli, J.B., and Haigis, M.C. (2018). The multifaceted contributions of mitochondria to cellular metabolism. Nature Cell Biology 20, 745–754.

Stroud, D.A., Formosa, L.E., Wijeyeratne, X.W., Nguyen, T.N., and Ryan, M.T. (2013). Gene knockout using transcription activator-like effector nucleases (TALENs) reveals that human NDUFA9 protein is essential for stabilizing the junction between membrane and matrix arms of complex I. The Journal of biological chemistry 288, 1685–1690.

Stroud, D.A., Surgenor, E.E., Formosa, L.E., Reljic, B., Frazier, A.E., Dibley, M.G., Osellame, L.D., Stait, T., Beilharz, T.H., Thorburn, D.R., et al. (2016). Accessory subunits are integral for assembly and function of human mitochondrial complex I. Nature 538, 123–126.

Thompson, K., Mai, N., Olahova, M., Scialo, F., Formosa, L.E., Stroud, D.A., Garrett, M., Lax, N.Z., Robertson, F.M., Jou, C., et al. (2018). OXA1L mutations cause mitochondrial encephalopathy and a combined oxidative phosphorylation defect. EMBO Molecular Medicine 10.

Tyanova, S., Temu, T., and Cox, J. (2016a). The MaxQuant computational platform for mass spectrometry-based shotgun proteomics. Nature Protocols 11, 2301–2319.

Tyanova, S., Temu, T., Sinitcyn, P., Carlson, A., Hein, M.Y., Geiger, T., Mann, M., and Cox, J. (2016b). The Perseus computational platform for comprehensive analysis of (prote)omics data. Nature Methods 13, 731–740.

Ugalde, C., Vogel, R., Huijbens, R., Van Den Heuvel, B., Smeitink, J., and Nijtmans, L. (2004). Human mitochondrial complex I assembles through the combination of evolutionary conserved modules: a framework to interpret complex I deficiencies. Human Molecular Genetics 13, 2461–2472.

Wallace, D., Singh, G., Lott, M., Hodge, J., Schurr, T., Lezza, A., Elsas, L., and Nikoskelainen, E. (1988). Mitochondrial DNA mutation associated with Leber's hereditary optic neuropathy. Science 242, 1427–1430.

Wittig, I., Braun, H.-P.P., and Schägger, H. (2006). Blue native PAGE. Nature Protocols 1, 418–428.

Zhu, J., Vinothkumar, K.R., and Hirst, J. (2016). Structure of mammalian respiratory complex I. Nature 536, 354–358.

Zurita Rendon, O., Silva Neiva, L., Sasarman, F., and Shoubridge, E.A. (2014). The arginine methyltransferase NDUFAF7 is essential for complex I assembly and early vertebrate embryogenesis. Human Molecular Genetics 23, 5159–5170.

