## Supplemental Figure 1 for "Optic Atrophy-associated TMEM126A is an assembly factor for the ND4-module of Mitochondrial Complex I"

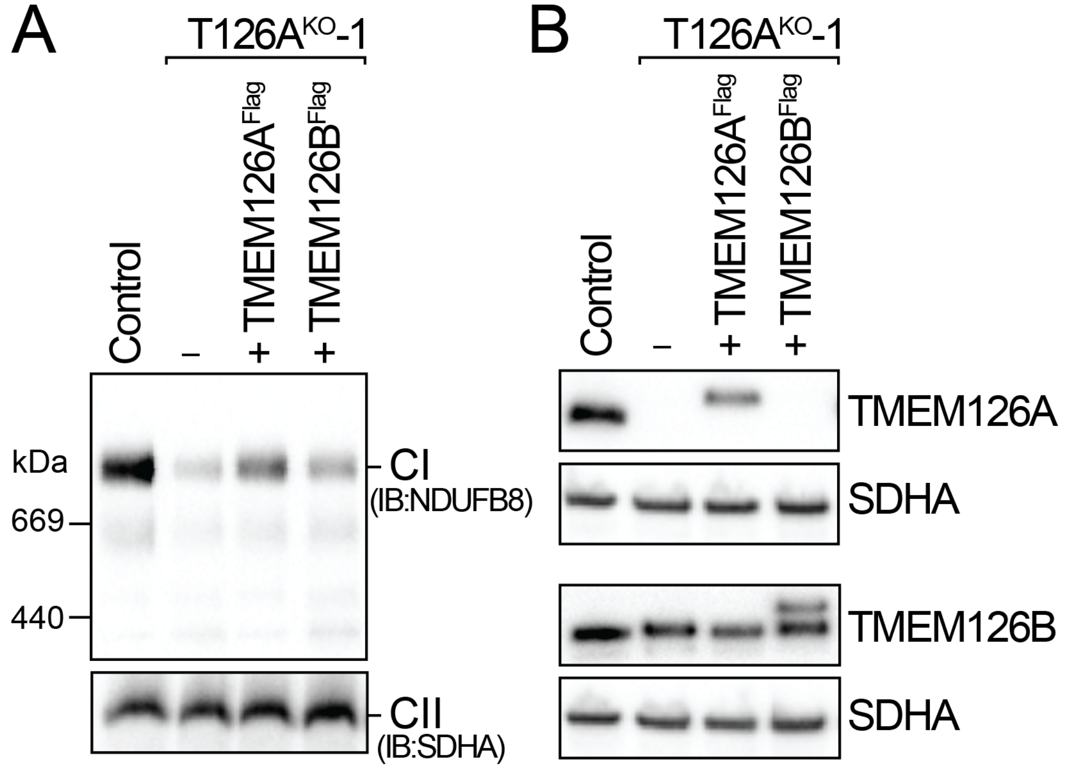


**Supplementary Figure 1: TMEM126A functions independently of TMEM126B.** Mitochondria isolated from control, T126A$^{KO}$-1 and T126A$^{KO}$-1 expressing TMEM126A$^{Flag}$ or TMEM126B$^{Flag}$ cells were analysed by **(A)** BN-PAGE after solubilization in 1% TX100 and immunoblotted for the complex I subunit NDUFB8 or **(B)** SDS-PAGE and immunoblotted for TMEM126A and TMEM126B. SDHA/CII was used as a loading control in all cases.
